# Extended access self-administration of methamphetamine is associated with age- and sex-dependent differences in drug taking behavior and recognition memory deficits in rats

**DOI:** 10.1101/850396

**Authors:** Sara R. Westbrook, Megan R. Dwyer, Laura R. Cortes, Joshua M. Gulley

## Abstract

Individuals who begin drug use during early adolescence experience more adverse consequences compared to those initiating later, especially if they are female. The mechanisms for these age and gender differences remain obscure, but studies in rodents suggest that psychostimulants may disrupt the normal ontogeny of dopamine and glutamate systems in the prefrontal cortex (PFC). Here, we studied Sprague-Dawley rats of both sexes who began methamphetamine (METH, i.v.) self-administration (SA) in adolescence (postnatal [P] day 41) or adulthood (P91). Rats received seven daily 2-h SA sessions with METH or saccharin as the reinforcer, followed by 14 daily long access (LgA; 6 h) sessions. After 7 and 14 days of abstinence, novel object (OR) or object-in-place (OiP) recognition was assessed. PFC and nucleus accumbens were collected 7 days after the final cognitive test and NMDA receptor subunits and dopamine D1 receptor expression was measured. We found that during LgA sessions, adolescent-onset rats escalated METH intake more rapidly than adult-onset rats, with adolescent-onset females earning the most infusions. Adolescent-onset rats exhibited modest deficits in OiP compared to adult-onset rats, but there was no sex difference in this effect and no groups differed in OR. We found no group differences in D_1_ and NMDA receptor expression, suggesting no long-lasting alteration of ontogenetic expression profiles. Our findings suggest that adolescent-onset drug use is more likely to lead to compulsive-like patterns of drug-taking and subsequent dysfunction of PFC-dependent cognition.

## Introduction

Most drug use begins during adolescence and those who initiate their use earlier in life appear to experience worse outcomes compared to those who start late in adolescence or during adulthood. For example, earlier onset of cocaine and other drug use has been associated with greater deficits in cognition in a battery of neuropsychological tests, a higher risk for psychosocial problems, and an increased risk for substance use disorder (SUD; Chen et al., 2009; Lopes et al., 2017; Poudel and Gautam, 2017). Female users also tend to experience worse outcomes, including a more rapid transition from initial to problematic drug use(Piazza et al., 1989; Haas and Peters, 2000; Brecht et al., 2004). Use of amphetamines, and especially the methylated derivative methamphetamine (METH), may be particularly problematic for these populations. Compared to males, females tend to initiate METH use earlier, are more sensitive to its acute behavioral and subjective effects, are more likely to have psychiatric problems associated with their drug use, and have worse treatment outcomes (Hser et al., 2005; Rawson et al., 2005; He et al., 2013; Mayo et al., 2019). In laboratory rats, females (Reichel et al., 2012a) and those beginning drug use during adolescence (Anker et al., 2012) develop compulsive-like METH seeking more readily as evidenced by greater and more rapid escalation of METH intake during extended access self-administration sessions. Together, these studies suggest that age-of-onset and sex may be factors that confer vulnerability to adverse outcomes of METH use, including a greater likelihood to develop compulsive METH-taking behavior and a heightened susceptibility to METH-induced cognitive dysfunction.

One hypothesized explanation for this heightened vulnerability is that drug use early in life may induce delays or other significant perturbations in normal brain development. METH, like nearly all other drugs of abuse, has potent effects in corticolimbic brain regions such as the nucleus accumbens and prefrontal cortex (PFC). These regions continue to reorganize and mature throughout late childhood and adolescence, with the PFC not reaching its mature, adult-like state until individuals are in their mid-to late twenties (Giedd, 2004). Moreover, this continued development is protracted compared to subcortical regions such as the nucleus accumbens (Mills et al., 2014). Studies using rodent models of adolescence have revealed a developmental shift in dopamine and glutamate signaling that occurs during adolescence. Specifically, dopamine D_1_ receptor (D_1_R) expression on PFC projections to the accumbens peaks during adolescence before pruning and relative decreases in expression occur as rats reach adulthood (Brenhouse et al., 2008). This D_1_R remodeling may precede the late adolescent emergence of GluN2B-containing NMDA receptor transmission in the PFC, which has been shown to be mediated by D_1_R signaling (Flores-Barrera et al., 2014). These ontogenetic changes are likely important mechanisms for developing adult-like cognition. In adults, intact D_1_R and NMDAR transmission in the medial prefrontal cortex (mPFC) are needed for certain forms of recognition memory. Pharmacological blockade of D_1_Rs in the mPFC impaired object-in-place (OiP) recognition memory, while sparing both novel object (NOR) and object location recognition memory (Savalli et al., 2015). Non-selective NMDAR blockade in the mPFC impairs OiP (Barker & Warburton, 2008). Although the role of different NMDAR subunits in OiP memory is not entirely clear, GluN2B function in the PFC has been implicated in working memory (Wang et al., 2013). Thus, drugs of abuse taken during adolescence may disrupt the ontogeny of D_1_R and/or GluN2B signaling in the developing PFC leading to greater deficits in OiP memory compared to adult-onset drug use, while sparing NOR memory.

Exposure to amphetamines during adolescence has been shown to influence the development of certain aspects of dopamine and glutamate signaling, and in turn cognitive functioning, though most of the published work to date has investigated age-of-onset or sex separately. In male rodents exposed to amphetamine non-contingently during adolescence, presynaptic sites on dopamine fiber inputs into the PFC are significantly reduced (Reynolds et al., 2015; Hoops et al., 2018), and dopamine-mediated inhibition of pyramidal cells in the PFC is significantly impaired at four- and twelve weeks after the last drug injection (Kang et al., 2016a; Paul et al., 2016). We have previously reported that these drug-induced neuroadaptations in the dopamine system are region specific, with reduced expression of D_1_Rs in the mPFC but no change in the NA after adolescent AMPH exposure (Kang et al., 2016b). Moreover, the drug-induced changes in pyramidal cell function are associated with significant disruptions in cognitive function, including impaired working memory (Sherrill et al., 2013), reduced impulse control (Hammerslag et al., 2014) and reductions in behavioral flexibility (Hankosky et al., 2013). A more recent study that examined the potential for age of exposure-dependent effects of METH on conditioned fear learning and extinction used only male rats and found adult-exposed animals to have deficits in extinction retrieval that were not apparent in their adolescent-exposed counterparts (Luikinga et al., 2019). The impact of adolescent amphetamine exposure on glutamate signaling has not been published to date, but a recent study demonstrated reduced expression of phosphorylated GluN2B in the infralimbic PFC after adolescent cocaine exposure in male rats (Caffino et al., 2018). Notably, most of the aforementioned studies employed non-contingent experimenter administered injections, and it is currently unknown whether psychostimulant-induced reductions in PFC D_1_Rs and GluN2B expression occur with contingent drug-taking during adolescence. Moreover, since most of these studies assessed adolescent drug exposure without including an adult-exposed comparison group, it is unclear whether observed adaptations are due to disruptions in the developmentally regulated processes specific to adolescence, or if they occur regardless of age of drug exposure.

The current study sought to address these gaps by using a methamphetamine (METH) self-administration paradigm to investigate the hypothesis that adolescent-onset METH-taking would disrupt the ontogenetic trajectories of D_1_R and GluN2B in the PFC in a sex-dependent fashion. To this end, we trained Sprague-Dawley rats of both sexes to self-administer METH or a non-drug reinforcer, saccharin, under short access (ShA) conditions beginning during adolescence or adulthood, followed by an extended period of long access (LgA) METH self-administration. One to two weeks following cessation of self-administration, rats were tested on object recognition memory tasks to assess METH-induced memory impairments. Seven days later, tissue was collected to assess NMDAR subunits and D_1_R protein expression in the PFC and NA. In line with previous work (Anker et al., 2012; Reichel et al., 2012a), we hypothesized that females and adolescent-onset rats would escalate their METH intake more rapidly during LgA compared to their male and adult-onset counterparts. Importantly, if METH-taking during adolescence indeed disrupted the ontogeny of D_1_R and GluN2B function in the PFC, we expected that adolescent-onset rats would experience greater deficits in PFC-dependent recognition memory and display greater reductions in D_1_R and GluN2B protein expression in the PFC, with no changes in the NA, as we found previously (Kang et al., 2016b). We further predicted that these METH-induced neuroadaptations may be more pronounced in females. Finally, we hypothesized that the effects would be specific to METH as a reinforcer, such that these patterns would not be evident in rats that self-administered the non-drug reinforcer, saccharin.

## Methods

### Subjects

Subjects were a total of 81 male and 84 female Sprague-Dawley rats that were born in-house on postnatal day (P) 1 from breeders originally obtained from Envigo (Indianapolis, IN, USA). Several rats were lost from the study due to issues with catheter patency (adolescent: males = 5, females = 3; adult: males = 3, females = 1), illness (adolescent: males = 3, females = 5), or other technical problems (adolescent females n = 2; adult: male n = 1; females = 1), yielding final subject totals of 96 males and 97 females. Rats were weaned on P22 and housed 2-3 same-sex animals per cage on a reversed 12-hour light/dark cycle (lights off at 0900). All behavioral training and testing occurred during the rats’ dark cycle. Food and water were available *ad libitum* throughout the study. Rats were weighed daily beginning on P25. Daily checks for physical markers to estimate puberty onset – preputial separation in males (Korenbrot et al., 1977) and vaginal opening in females (Castellano et al., 2011) – began on P30 and continued until all rats were in puberty. Assignment to groups based on age-of-onset (adolescent or adult) and reinforcer (METH or saccharin) was counterbalanced across litters. All procedures were approved by the Institutional Animal Care and Use Committee at the University of Illinois and followed the National Research Council’s Guide for the Care and Use of Laboratory Animals.

### Self-administration

The timeline for experimental procedures is shown in Figure 1. Intravenous catheterization surgery was performed as previously described (Hankosky et al., 2018) on P32 (± 2 days) or P82 (± 7 days) for adolescent-onset and adult-onset groups, respectively. Rats were given at least 5 days to recover from surgery prior to beginning daily self-administration sessions. Antibiotic (1.1% trimethoprim sulfa; Midwest Veterinary Supply) was administered via the drinking water starting one day before surgery and continuing throughout the experiment. Catheters were flushed daily with 0.1 ml of 50 U/ml heparinized saline. Catheter patency was assessed once following surgery recovery, and as needed if patency loss was suspected, by infusion of ≤ 0.1 ml of a 15% ketamine (100 mg/ml) and 15% midazolam (5 mg/ml) solution. Loss of muscle tone shortly after infusion was taken as a positive indicator of catheter patency.

**Figure 1.**
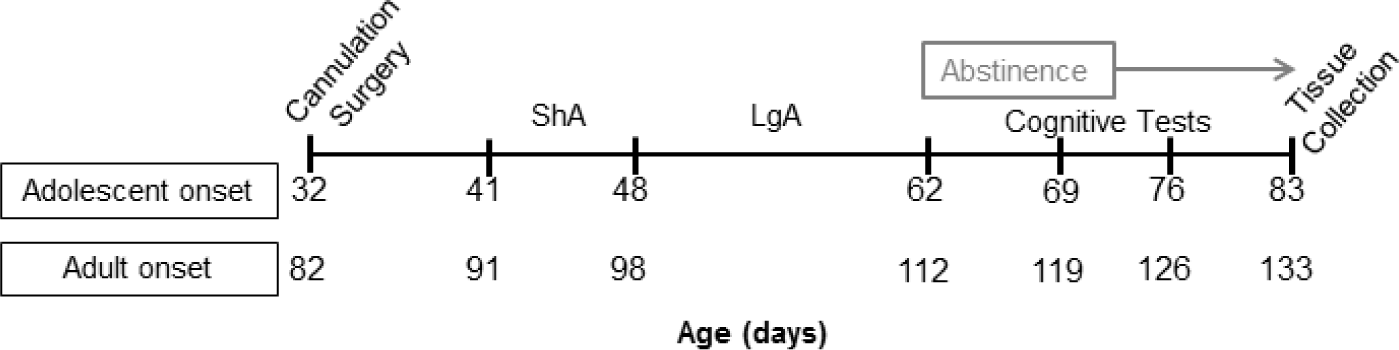
Experimental timeline for rats that underwent self-administration under short access (ShA) and long access (LgA) conditions. Cognitive tests included novel object recognition and object-in-place recognition tasks after 7 and 14 days of abstinence with task order counterbalanced. mPFC and NA were collected after 21 days of abstinence.

On P40 (± 2 days) or P90 (± 7 days), rats received one 90-min habituation session in the operant chambers with the nosepoke ports blocked. The following day rats were trained in 7 daily, ShA sessions (2 h duration) to nosepoke into one of two recessed ports in order to receive an intravenous infusion of METH (0.1 mg/kg/infusion) or a liquid dipper cup (0.06 or 0.08 mL) of 0.1% saccharin. The duration of the METH infusion was varied between 2 and 4 sec in order to maintain the unit dose of 0.1 mg/kg/infusion across days using rats’ daily body weights. Nosepoke responses into the active nosepoke port resulted in reinforcement on an FR1 schedule, whereas responses into the inactive nosepoke port had no programmed consequences. Each reinforcer delivery coincided with illumination of the cue light above the active port and the presentation of a tone for 4 seconds. Deliveries were followed by a 20-s timeout period during which the nosepoke ports were illuminated with a red light and responses into the ports had no programmed consequences. Following ShA sessions, rats received 14 daily LgA sessions that were 6 h in duration but otherwise identical to ShA sessions. In order to minimize risk of overdose, rats responding for METH could earn a maximum of 120 infusions per LgA session.

### Recognition Memory

Rats with a history of METH or saccharin self-administration, as well as naïve littermates (aged P93 at first test) from the final birth cohort of rats, were tested in a counterbalanced order on novel object recognition (NOR; Fig. 5A) and object-in-place recognition (OiP; Fig. 6A) memory tasks 7 and 14 days after self-administration sessions ended. Our procedures for these tests were adapted from our previous studies (Westbrook et al., 2014) and others (Barker et al., 2007; Reichel et al., 2012a). Each task consisted of three 10-min habituation sessions, a 5-min study phase, a 90-min delay, and a 3-min test phase. No objects were present during habituation sessions. Sessions were run over two days with one morning session followed by another session 5 h later. For the NOR task, two identical objects were present during the study phase and one of these objects was replaced with a novel object during the test phase. For the OiP task, four different objects were present during the study phase. During the test phase, the locations of two of the objects were switched with each other. All objects were of similar size and could be sanitized, but differed in shape, color, and texture. Testing occurred in open-field arenas (41 x 41 x 41 cm) with clear acrylic walls. Visual cues—pink and yellow plaid design, white paper with a black X, and blue and red vertical stripes—were adhered to the outside of the west, north, and east walls, respectively, to act as proximal spatial cues. Arenas and objects were wiped clean with 70% ethanol between all sessions in order to eliminate odor cues.

Three weeks after the last self-administration session (one week after the final recognition task), rats were anesthetized with 195 mg/kg pentobarbital and transcardially perfused with 30-40 mL of ice-cold saline. Brains were removed, chilled in saline for approximately 1 min prior to sectioning using an ice-cold metal brain matrix. Bilateral samples of the vmPFC and NA were extracted using a 2.0 mm diameter punch from 1 mm coronal slices. Prelimbic and infralimbic vmPFC were combined in accordance with our previous work that showed no differential effect of AMPH exposure on D_1_R function in these subregions (Kang et al., 2016a,b).

In a subset of 11 rats per group, D_1_R (1:2000, ab20066, Abcam; Kang et al., 2016b), GluN2B (1:1000, 4207, Cell Signaling, Hafenbreidel et al., 2014), and GluN1 (1:1000, ab52177, Abcam; Sun et al., 2013) protein expression was measured via Western blot as described in Kang et al. (2016b). Although we observed multiple bands using this D_1_R antibody, which is not uncommon for D_1_R antibodies in general, we used the band at ∼50 kDa for our analyses since we and others have previously used this approach for the current D_1_R antibody (De Santis et al., 2016; Kang et al., 2016b; Zhu et al., 2017). Briefly, brain tissue was homogenized in lysis buffer, centrifuged, and protein concentration estimated in the supernatant using Precision Red Advanced Protein Assay (Cytoskeleton, CA). Samples were prepared at 25 µg protein/well with 2X laemmli loading buffer and 2-mercaptoethanol and run on pre-cast 4-15% gels (Biorad). Proteins were wet transferred onto PVDF membranes, blocked with 5% non-fat milk in TBST, and incubated overnight with one of the primary antibodies listed above. The next day, membranes were washed, incubated with secondary antibody, washed, and incubated with horseradish substrate prior to imaging. Following imaging, membranes were washed and stripped for re-probing with anti-GAPDH (1:1000, ab9484, Abcam; Kang et al., 2016b), a house-keeping protein, used as a loading control. A homogenate of naïve adult female rats was used to normalize across gels. A single gel consisted of one well each of the naïve adult female homogenate, adolescent-onset METH male, adolescent-onset METH female, adolescent-onset saccharin male, adolescent-onset saccharin female, adult-onset METH male, adult-onset METH female, adult-onset saccharin male, and adult-onset saccharin female.

## Data Analysis

### Body weights

Starting on P25, body weight was measured for all rats. For each sex, body weights from P25 – P74 were analyzed with separate two-way mixed ANOVAs for adolescent-onset rats. Separate two-way ANOVAs for each sex were conducted on body weights of adult-onset rats from P75-120. In these analyses, the between-subjects factor was reinforcer and the within-subjects factor was postnatal day.

### Self-administration

Reinforcers earned were measured for each ShA and LgA session. Between-subjects factors included sex and age-of-onset, while the within-subjects factor was session. All analyses were conducted separately for rats that self-administered METH and saccharin with reinforcer intake (mg/kg or L/kg) as the dependent measure. Separate three-way mixed ANOVAs (sex x age-of-onset x session) were used to assess reinforcer intake during ShA (days 1-7) and LgA (days 8-21) sessions. Escalation during LgA sessions was assessed via separate one-way repeated measures ANOVAs within each sex/age-of-onset group with Tukey post-hoc comparisons of reinforcer intake in each LgA session (days 9-21) compared to the first LgA session (day 8). These analyses were conducted using total session intake for the entire session as well as reinforcer intake during the first hour of LgA sessions. Cumulative METH (mg/kg) and saccharin (ml/kg) intake was calculated for all self-administration sessions and analyzed using separate two-way (sex x age-of-onset) ANOVAs for each reinforcer. Tukey post-hoc tests were conducted where appropriate for all analyses.

### Recognition memory tasks

Time spent exploring the objects during the study and test phase was measured. Exploration ratios were calculated as time spent exploring the novel object or switched objects divided by the total time spent exploring all objects. An exploration ratio of 0.5 indicates chance performance, i.e. equal exploration of novel and familiar objects. Separate analyses were conducted for each recognition task. Two-way factorial ANOVAs with age-of-onset and sex as factors for rats with a self-administration history (or only sex as a factor for naïve control rats) were conducted on total exploration time (sec) during the 5-min study phase to assess potential baseline differences in exploratory activity. Total exploration ratios for the 3-min test phase were analyzed separately for rats based on self-administration history using two-way factorial ANOVAs with age-of-onset and sex as factors (only sex as a factor for naïve rats). We also analyzed test phase exploration ratios for each min of the test phase as previous studies (Dix and Aggleton, 1999; Jablonski et al., 2013) have reported that the first minute may be a more sensitive measure since this is when the novel change is the most salient. In order to assess these changes in exploration as the test phase progressed, three-way mixed ANOVAs with age-of-onset, sex, and minute (within-subjects) as factors were conducted on exploration ratios. In order to compare rats with a self-administration history to naïve control rats, two-way ANOVA with group (i.e., naïve, adolescent-onset METH, adult-onset METH, adolescent-onset saccharin, and adult-onset saccharin) and sex as factors was conducted on total exploration ratios for the test phase. Additionally, three-way ANOVA with group, sex, and minute (within-subjects) as factors was used to compare the time course of exploration ratios across the test phase. Lastly, test phase total exploration ratios and individual minute exploration ratios for each group were compared to chance performance (0.5) using one-sample t-tests as this is a common statistical convention within the recognition task literature to determine whether each group demonstrated significant preference for the novel change (Dix and Aggleton, 1999; Westbrook et al., 2014; Ramsaran et al., 2016). To account for multiple testing, we used the false discovery rate adjustment for p-values (less than 5%; Benjamini and Hochberg, 1995).

### Protein expression

Optical density of the protein bands of interest (D_1_R: ∼50 kDa, GluN2B: ∼180 kDa; GluN1: ∼120 kDa) was divided by the density of GAPDH (∼39 kDa). Data were normalized across gels by dividing the protein expression for each group by the expression of naïve adult female homogenate (control). The relative intensity of the bands was calculated using the following equation: [(Protein of interest/GADPH)/(Control protein of interest/Control GAPDH)]. Relative intensities for each protein of interest (D_1_R, GluN2B, and GluN1) was analyzed separately for rats with a history of METH or saccharin self-administration using two-way ANOVAs with age-of-onset and sex as factors for each brain region.

## Results

### Self-administration

#### Body Weight

Separate two-way ANOVAs by sex and age-of-onset revealed significant main effects of postnatal day [Adolescent-onset: females F(49,1262)=90.08, p<0.0001; Adult-onset: males F(45,1211)=9.12, p<0.0001, females F(45,1435)=5.27, p<0.0001], which was due to the expected weight gain as Sprague-Dawley rats age under ad libitum access to food (Fig. 2). In adolescent-onset males, two-way ANOVA revealed a significant reinforcer by postnatal day interaction [F(49,1167)=1.58, *p*=0.0076, reflecting that male rats that self-administered METH during adolescence had modest, but significant, reductions in weight gain compared to their saccharin self-administering counterparts starting on P55. These body weight differences were no longer statistically significant beginning on P67 (∼1 week after self-administration ended). Notably, this weight gain suppressing effect of METH was only evident in adolescent-onset males.

**Figure 2.**
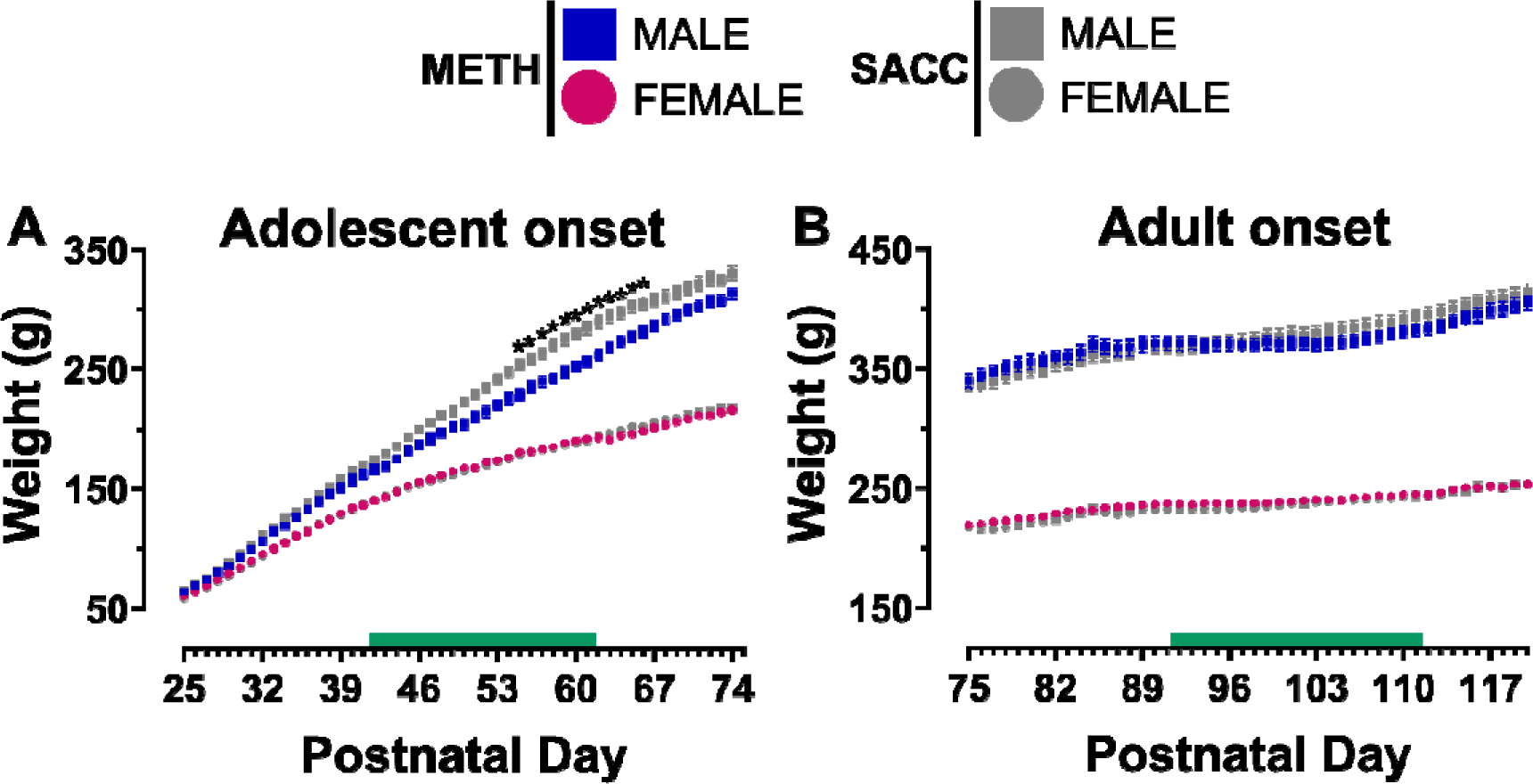
Body weight (g) across postnatal days (P) 25-74 for adolescent-onset rats (**A**) and P75-120 for adult-onset rats (**B**). The green bar indicates the ages when rats underwent METH or saccharin (SACC) self-administration sessions. In this and all subsequent figures, data expressed as mean ± SEM. * *p* < 0.05 vs. males that self-administered METH on the noted days.

#### Short Access

In METH self-administering rats, three-way ANOVA of intake during ShA sessions revealed only a significant main effect of session [F(6,55)=10.14, *p*<0.0001], with METH intake higher on session 1 compared to all other ShA sessions (all *p*’s<0.0002). After session 1, ShA intake of METH remained stable and did not differ statistically (Fig. 3A). Three-way ANOVA of saccharin intake (Fig. 4A) across ShA sessions showed significant main effects of sex [F(1,54)=50.57, *p*<0.0001] and session [F(6,54)=42.46, *p*<0.0001], as well as a significant sex by session interactions [F(6,54)=8.12, *p*<0.0001]. During ShA sessions, females increase their saccharin intake from session 1 to 2 (*p*<0.0001) before their intake plateaued, whereas males do not exhibit any significant stepwise changes in saccharin intake across ShA sessions. Moreover, females earn significantly more saccharin than males for all ShA sessions.

**Figure 3.**
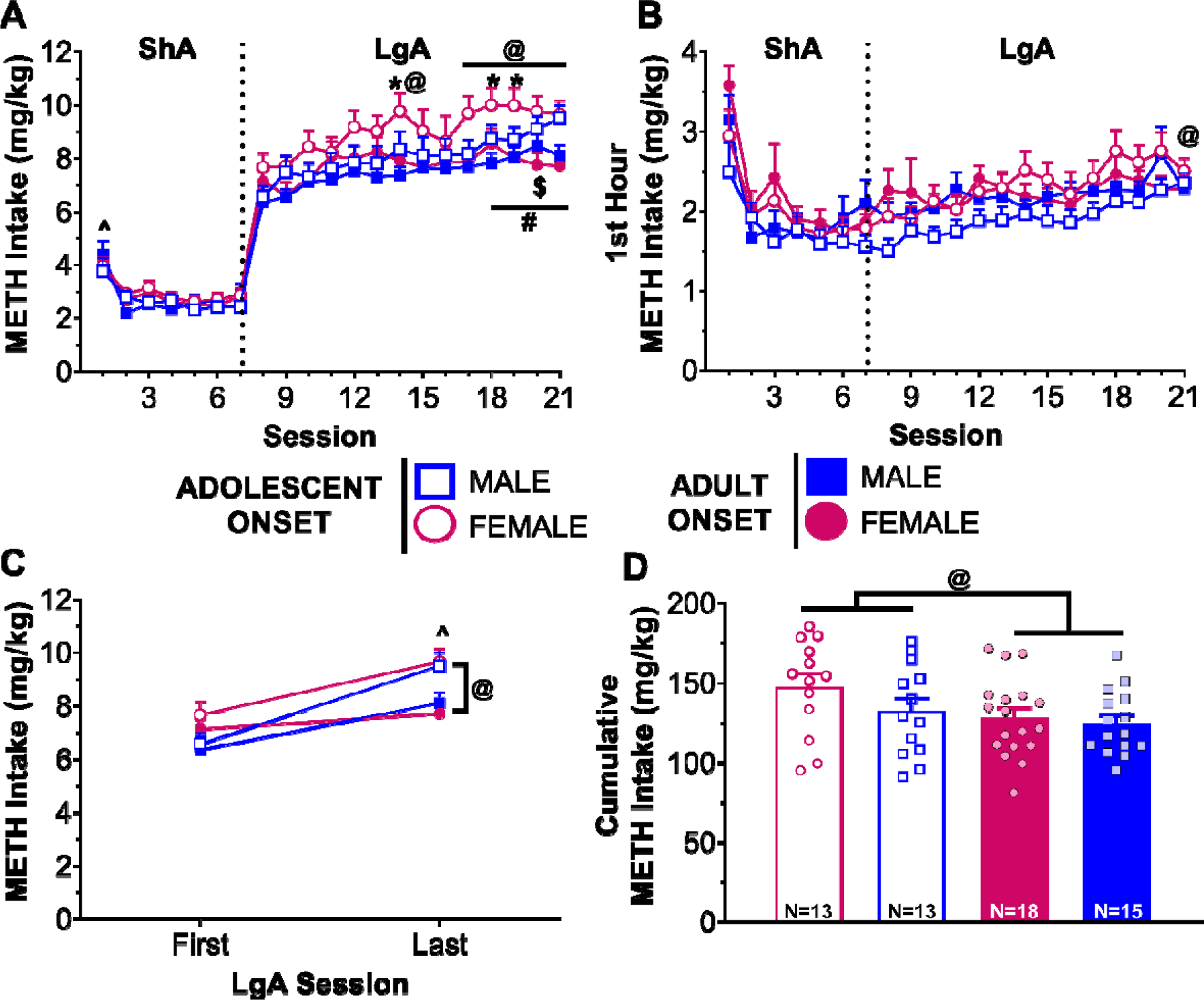
Adolescent- and adult-onset METH self-administration in rats of both sexes. (**A**) Daily METH intake (mg/kg) across 7 days of short access (ShA) and 14 days of long access (LgA) self-administration. ^*p* < 0.05 vs. all other ShA sessions. Session intake vs. first LgA session: * *p* < 0.05 for females, ^#^ *p* < 0.01 for males, ^@^ *p* < 0.05 for adolescent-onset rats, ^$^ *p* < 0.05 for adult-onset rats. (**B**) Daily METH intake (mg/kg) during the first hour of each self-administration session. ^@^ *p* < 0.05 vs. first LgA session for adolescent-onset rats. (**C**) METH intake (mg/kg) on the first and last LgA session. ^@^ *p* < 0.05 age within session, ^*p* < 0.05 vs. first LgA session. (**D**) Cumulative METH intake (mg/kg) across all self-administration sessions. Each dot represents one individual rat. ^@^ *p* < 0.05 age effect.

**Figure 4.**
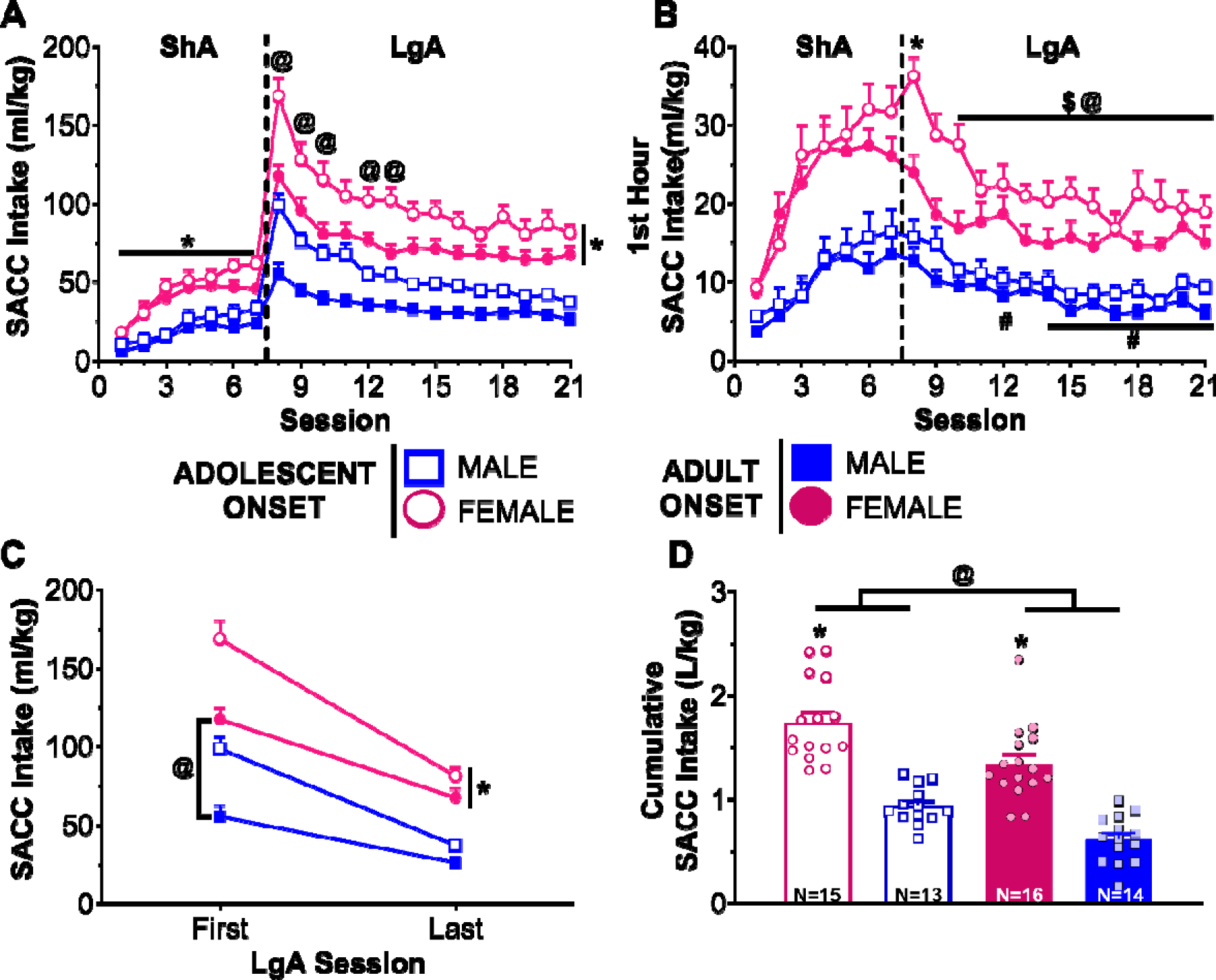
Adolescent- and adult-onset saccharin self-administration in rats of both sexes. (**A**) Daily saccharin (SACC) intake (ml/kg) across 7 days of short access (ShA) and 14 days of long access (LgA) self-administration. * *p* < 0.05 vs. males, ^@^ *p* < 0.05 adolescent-onset vs. adult-onset. (**B**) Daily SACC intake (ml/kg) during the first hour of each self-administration session. * *p* < 0.05 for females vs. all other LgA sessions, Session intake vs. first LgA session: ^#^ *p* < 0.01 for males, ^@^ *p* < 0.05 for adolescent-onset rats, ^$^ *p* < 0.05 for adult-onset rats. (**C**) SACC intake (ml/kg) on the first and last LgA session. * *p* < 0.05 vs. males, ^@^ *p* < 0.05 vs. adolescent-onset. (**D**) Cumulative SACC intake (L/kg) across all self-administration sessions. Each dot represents one individual rat. * *p* < 0.05 vs. males, ^@^ *p* < 0.05 age effect.

#### Long Access

During LgA sessions (Fig. 3A), three-way ANOVA conducted on METH intake showed significant age by session [F(13,55)=3.88, *p*=0.0002] and sex by session [F(13,55)=2.79, *p*=0.0040] interactions. Post-hoc analyses of the age by session interaction indicated that adolescents increase METH intake relative to session 1 starting on session 14 (*p*=0.0022), whereas adults did not exhibit this increase until session 20 (*p*=0.0421). Females begin increasing their METH intake earlier than males with females at session 14 (*p*=0.0425) and males at session 19 (*p=*0.0038). Three-way ANOVA conducted on saccharin intake across LgA sessions (Fig. 4A) revealed significant age by session [F(13,54)=3.42, *p*=0.0007] and sex by session [F(13,54)=2.32, *p*=0.0156] interactions. In addition to adolescent-onset rats having higher saccharin intake relative to adult-onset rats across all sessions (all *p*’s<0.041), adolescent-onset rats also reduced their saccharin intake relative to session 8 earlier than adult-onset rats (adolescent-onset: session 9, *p*<0.0001; adult-onset: session 10, *p*=0.0208). A similar pattern was seen for females compared to males. Females maintained higher saccharin intake relative to males (all *p*’s<0.001), but they reduced their intake more rapidly than males (females: session 9, *p*<0.0001; males: session 12, *p*<0.0001).

Three-way ANOVA using only the first hour of METH intake during LgA sessions (Fig. 3B) revealed significant age by session [F(13,55)=1.92, *p*=0.0482] and sex by session [F(13,55)=2.17, *p*=0.0233] interactions. Post-hoc analyses of the sex by session interaction showed no significant differences in first hour METH intake from session 8, whereas the age by session interaction indicated that adolescent-onset rats increased their first hour METH intake from the first to the last LgA session (*p*=0.0125). Analyses of the first hour saccharin intake (Fig. 4B) showed significant age by session [F(13,702)=2.29, *p*=0.0057] and sex by session [F(13,702)=2.30, *p*=0.0055] interactions. Both age groups significantly reduced their first hour saccharin intake starting on session 10 (adolescent-onset: *p*<0.0001, adult-onset: *p*=0.0077). In addition, adolescent-onset rats had higher saccharin intake for the first 2 sessions compared to adult-onset rats (all *p*’s<0.028). Females reduce their first hour saccharin intake starting on session 9 (*p*<0.0001), while males began reducing on session 12 (*p*=0.00334). Throughout the LgA sessions, females maintained higher first hour intake of saccharin than males (all *p*’s<0.0033).

Escalation was further assessed using a three-way ANOVA comparing METH intake on only the first and last LgA sessions (Fig. 3C). This test revealed a significant age by session interaction [F(1,54)=5.92, *p*=0.0183]. Initially, adolescent-onset and adult-onset rats’ METH intake did not differ statistically on the first LgA session. However, METH intake increased in both age groups from the first to the last LgA session (adolescent-onset: *p*<0.0001; adult-onset: *p*=0.0069), with adolescent-onset rats taking significantly more METH by the last LgA session compared to adults (*p*=0.0074). A similar analysis of saccharin intake (Fig. 4C) revealed age by session [F(1,54)=21.18, *p*<0.0001] and sex by session [F(1,54)=9.40, *p*=0.0034] interactions. Both age groups reduced their saccharin intake from the first to the last LgA session (both *p*’s<0.0001), but adolescent-onset rats took more saccharin than their adult-onset counterparts during both sessions (first LgA: *p*<0.0001, last LgA: *p*=0.0406). Similarly, both sexes reduced their intake (both *p*’s<0.0001) with females maintaining higher saccharin intake than males (both *p*’s<0.0001).

Lastly, cumulative METH intake was calculated across all self-administration sessions and is illustrated in Fig. 3D. Two-way ANOVA indicated a significant main effect of age [F(1,55)=4.03, *p*=0.0496], with adolescent-onset rats having greater total intake than adult-onset rats, regardless of sex. Cumulative saccharin intake (Fig. 4D) significantly differed by age [F(1,54)=19.00, *p*<0.0001] and sex [F(1,54)=86.41, *p*<0.0001], but there was not a significant interaction. Adolescent-onset and female rats had higher cumulative saccharin intake than adult-onset and male rats, respectively.

### Recognition memory

#### NOR task

During the study phase, total object exploration time was assessed using separate two-way ANOVAs based on self-administration history (METH, saccharin, or naïve). No significant main effects or interactions were revealed for rats regardless of self-administration history. This indicates that there were no baseline differences in exploration times during the study phase that might contribute to differences seen during the test phase.

Separate two-way ANOVAs conducted on total exploration ratios during the test phase (Fig. 5) for rats that previously self-administered METH or saccharin revealed no significant main effects or interactions. A similar lack of significant effects was found for naive control rats. Time course analyses of exploration ratios across the three-min test phase revealed no significant main effects or interactions in rats with a history of METH self-administration, whereas in rats with a history of saccharin self-administration there was a significant main effect of min [F(2,54)=13.93, *p*<0.0001] and an age by min interaction [F(2,54)=4.38, *p*=0.0173]. Rats in the adolescent-onset group had a significantly higher exploration ratio in min 1 compared to min 3 (*p*<0.0001). No significant main effects or interactions were reported from the two-way ANOVA comparing total exploration ratios for rats with a self-administration history to naïve control rats. Time course analyses including naïve control rats indicated a significant main effect of min [F(2,131)=15.06, *p*<0.0001], such that exploration ratios in the first min were significantly greater than the second and third min (*p*’s<0.001).

**Figure 5.**
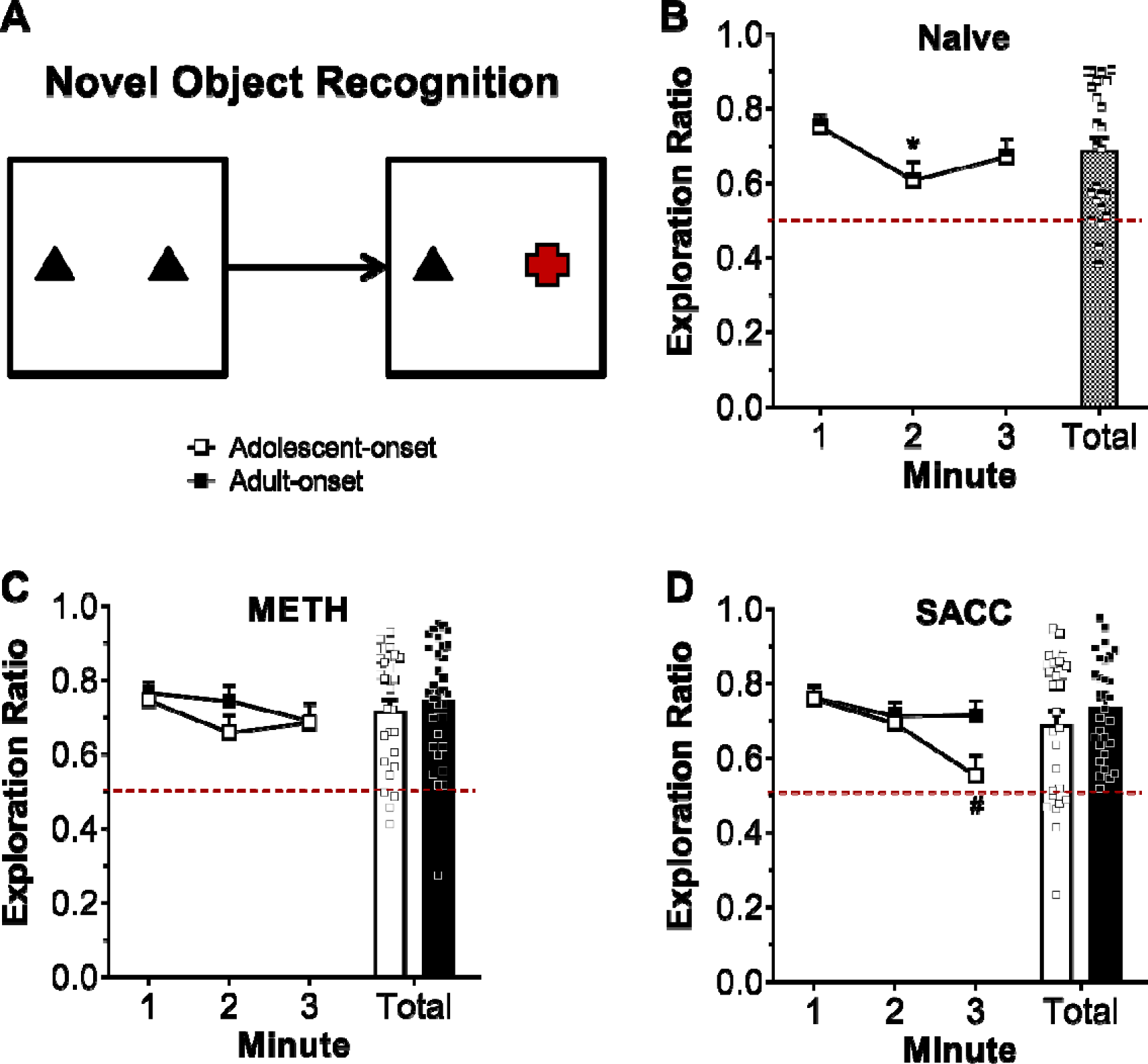
Novel object recognition memory assessed during abstinence from METH or saccharin self-administration. Schematic of the novel object recognition task with red object indicating the novel target (**A**). Exploration ratios during the test phase for naïve rats (**B**) or rats with a history of METH (**C**) or SACC (**D**) self-administration. Exploration ratios were calculated as time_novel_ / (time_novel_ + time_familiar_). Dotted red line indicates chance performance (0.5). We found no significant sex difference, so data are presented collapsed across sex. * *p* < 0.05 vs. min 1, ^#^ *p* < 0.001 vs. min 1 within adolescent-onset.

Since there was no evidence of sex differences in any of the NOR task analyses, groups were collapsed across sex for the one-sample t-tests of test phase exploration ratios against chance performance (0.5). For total exploration ratios (Table S1), all groups demonstrated preference for the novel object that was significantly greater than chance (all FDR-corrected *p*’s ≤ 0.0003). Rats also displayed significant novelty preference in the individual minute analyses (Table S2).

#### Object-in-place (OiP) task

Total object exploration time was examined during the study phase with separate two-way ANOVAs by self-administration history. The ANOVAs for rats from each group failed to indicate any significant main effects or interactions. Similar to the findings in the NOR task, there were no baseline differences in exploration during the study phase.

For the test phase (Fig. 6), two-way ANOVAs were conducted on total exploration ratios. These analyses revealed a significant main effect of age-of-onset [F(1,55)=4.79, *p*=0.0329] for rats with METH self-administration history and no significant main effects or interactions for rats that responded for saccharin. Rats that self-administered METH beginning during adolescence had lower test phase exploration ratios compared to their adult-onset counterparts, regardless of sex. Time course analyses including test phase minute as a factor indicated significant main effects of age-of-onset [F(1,55)=4.49, *p*=0.0386] and minute [F(2,55)=3.51, *p*=0.0369] for rats with a history of METH self-administration and no significant main effects or interaction for rats with a history of saccharin self-administration. Two-way ANOVA comparing total exploration ratios for rats with a self-administration history to naïve control rats revealed no significant main effects or interactions. However, time course analyses including naïve control rats indicated a significant group by min interaction [F(8,131)=2.90, *p*=0.0052]. Exploration ratios for naïve control rats significantly decreased from the first min to the third minute (*p*=0.0447), and this effect was absent in the groups with a history of self-administration.

**Figure 6.**
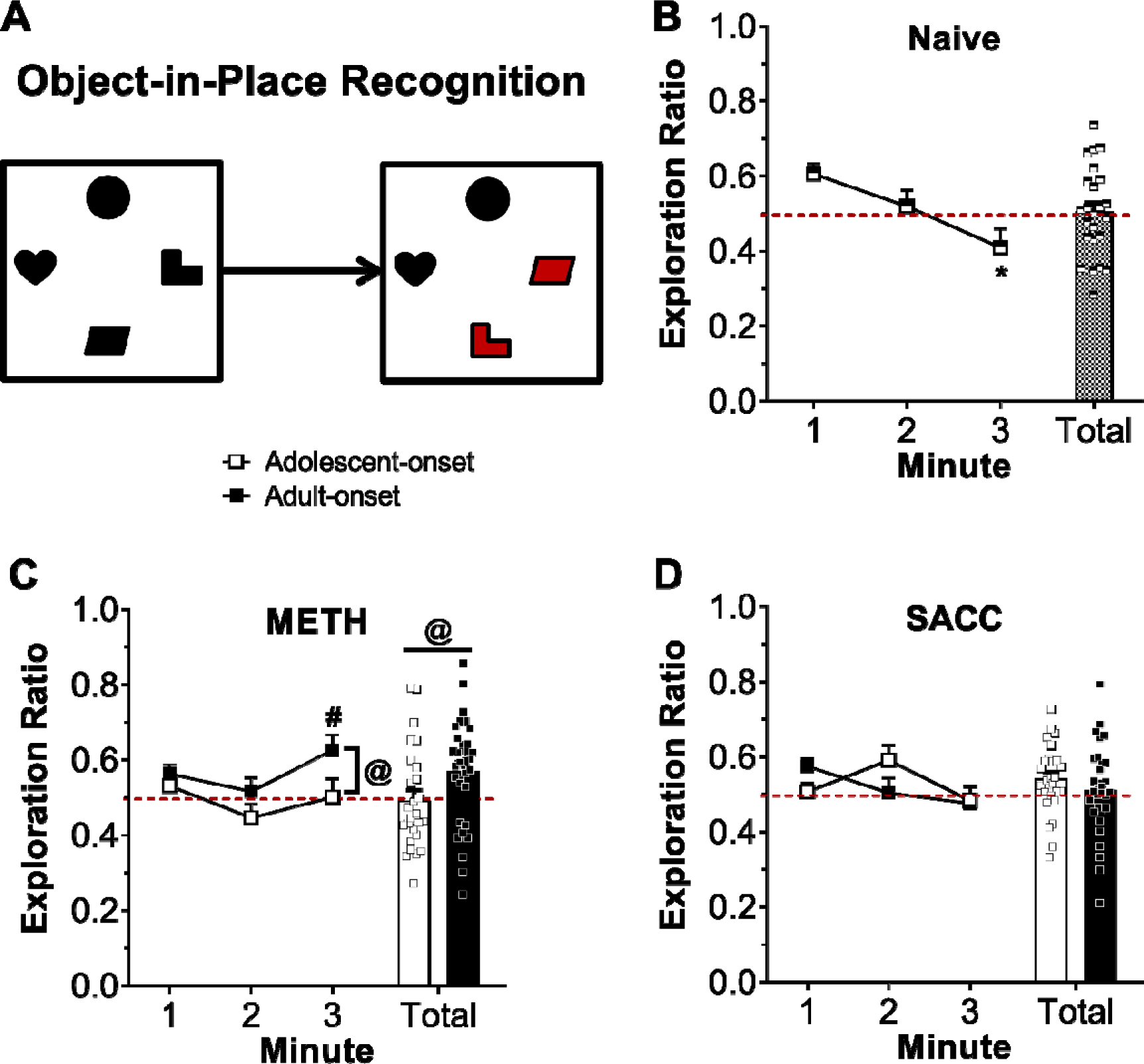
Object-in-place recognition memory assessed during abstinence from METH or saccharin self-administration. Schematic of the object-in-place recognition task with red objects indicating the novel targets (**A**). Exploration ratios during the test phase for naïve rats (**B**) or rats with a history of METH (**C**) or SACC (**D**) self-administration. Exploration ratios were calculated as time_novel_ / (time_novel_ + time_familiar_). Dotted red line indicates chance performance (0.5). We found no significant sex difference, so data are presented collapsed across sex. * *p* < 0.01 vs. min 1, ^#^ *p* < 0.05 vs. min 2 within adult-onset, ^@^ *p* < 0.05 age-of-onset effect collapsed across min.

Groups were collapsed across sex for the one-sample t-tests of test phase exploration ratios against chance performance (0.5) due to the lack of significant effects of sex in the object-in-place recognition task analyses. Not all groups demonstrated preference for the novel targets that was significantly greater than chance performance when examining the total exploration ratios (Table S1). However, the individual minute exploration ratio analyses (Table S2) demonstrated that all groups displayed significant preference for the novel targets during the first minute of the test phase, except for rats with adolescent-onset METH or saccharin history. Moreover, rats that self-administered METH starting during adolescence did not display significant preference for the novel object during any minute of the test phase. In contrast, rats with a history of adolescent-onset saccharin self-administration demonstrated significant preference for the switched objects during minute 2 of the test phase [t(27)=5.06, FDR-corrected *p*=0.045], indicating that they were able to recognize the novel change.

#### Receptor expression

A randomly selected subset of rats (n=11/group) were used for Western blot analyses of receptor protein expression. One rat’s NA sample (a male from the adolescent-onset METH group) was lost due to an error in sample preparation. For data from the remaining samples, separate two-way ANOVAs were conducted on the relative intensity of each protein band of interest (D_1_R, GluN1, GluN2B) for each brain region (PFC and NA) and by self-administration history. We observed no significant main effects or interactions for any of these measures (Fig. 7).

**Figure 7.**
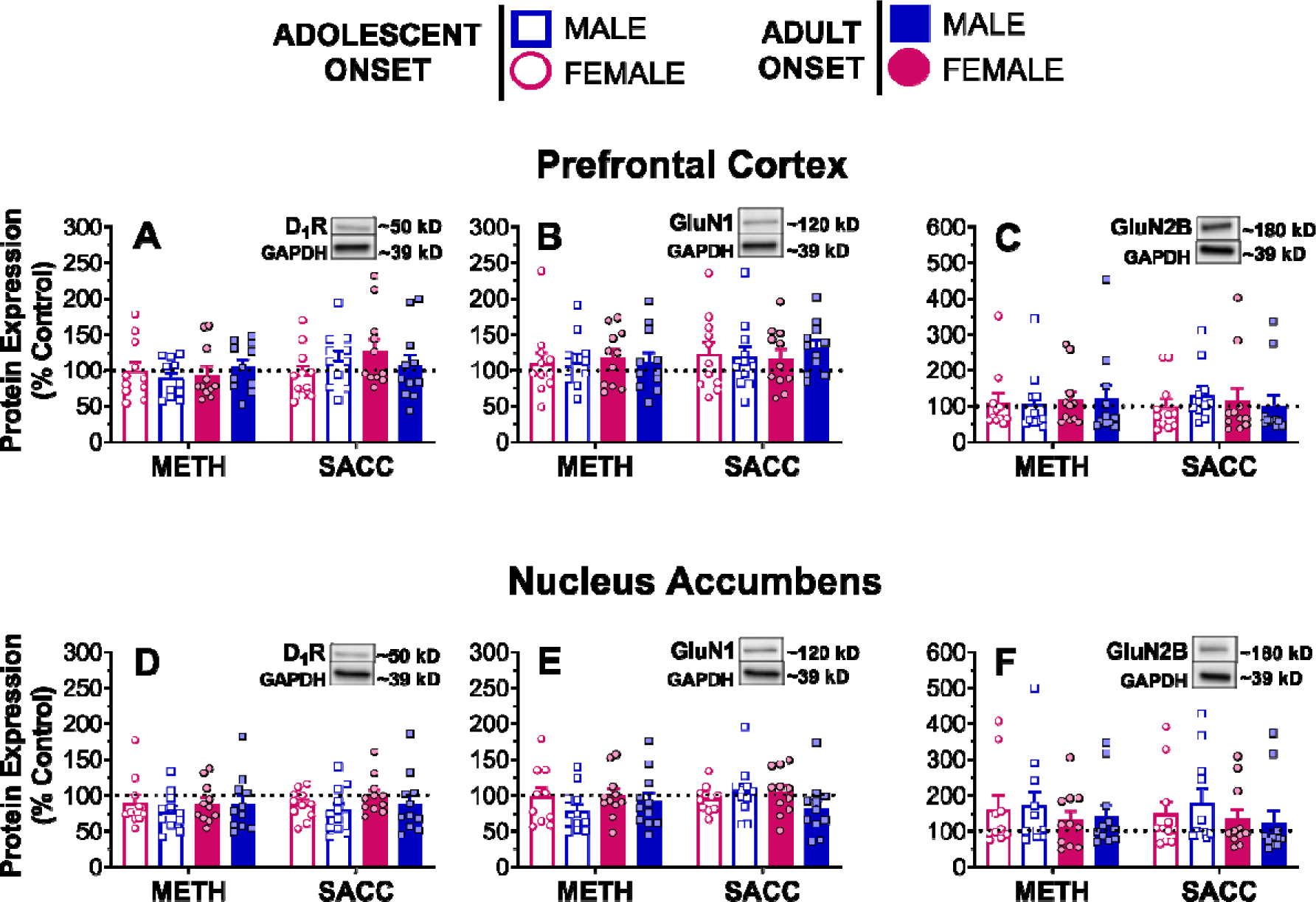
Dopamine D1 receptor and NMDA receptor subunits protein expression as a percent of naïve adult female homogenate (control). Tissue was collected 21 days after the last self-administration session. Bands are shown for the control homogenate. Each dot represents one individual rat.

## Discussion

Adolescence is characterized by considerable development of D_1_R and GluN2B neurotransmission in the PFC (Andersen et al., 2000; Brenhouse et al., 2008; Flores-Barrera et al., 2014), which may constitute a window of vulnerability to drug-induced neuroadaptations (Gould, 2010; Gulley and Juraska, 2013). This vulnerability may partly explain why adolescent-onset drug users suffer worse outcomes of their drug use compared to adult-onset users. We tested this hypothesis in the current study by investigating PFC-dependent cognition and receptor expression in rats with a history of METH or saccharin self-administration. We found that adolescent-onset and female rats escalated METH intake more rapidly and maintained higher saccharin intake than their adult-onset and male counterparts. Regardless of sex, rats that initiated METH self-administration during adolescence were modestly impaired in OiP recognition memory compared to adult-onset rats. Despite the age-of-onset difference in PFC-dependent cognition, D_1_R and NMDAR subunit protein expression was not altered three weeks after the last drug-taking session. Our findings suggest that METH self-administered during adolescence impairs PFC-dependent cognition; however, METH does not appear to induce long-lasting disruptions of D_1_R or GluN2B developmental expression trajectories in the PFC.

Consistent with previous reports from our lab (Hankosky et al., 2018) and others (Ahmed and Koob, 1998; Kitamura et al., 2006), METH intake was relatively stable in both adolescent- and adult-onset groups when the drug was available under ShA conditions. During ShA, METH intake was significantly higher on the first session compared to the rest of the ShA sessions. During habituation, the nosepoke ports were blocked. This practice makes the nosepoke ports more salient when the rats have access to them during the first self-administration, which likely contributes to the increased intake during this session. After the first ShA session, METH intake for all groups stabilized. Interestingly, group differences emerged across sessions when rats could self-administer METH under LgA conditions. Consistent with a previous report in males (Anker et al., 2012), adolescent-onset rats escalated more rapidly than adult-onset rats. Also in line with previous work (Reichel et al., 2012a), females began increasing their METH intake earlier than males, although this effect due largely to adolescent-onset females. One explanation for the age-of-onset difference in escalation is that adolescent rats may be less sensitive to METH and therefore need to take more drug to achieve a similar effect. We have previously reported that adolescent-onset rats are less sensitive to reinforcement with a low dose of METH (0.02 mg/kg/inf), and adolescent-onset rats reach lower breakpoints in progressive ratio (PR) compared to adult-onset rats (Hankosky et al., 2018). In the present study, we used a relatively high unit dose of METH (0.1 mg/kg/inf) and we did not see any age-of-onset differences during ShA sessions. Also inconsistent with a reduced sensitivity in adolescence is that locomotor sensitization to psychostimulants is often enhanced or equal in adolescents compared to adults (Adriani et al., 1998; Mathews and McCormick, 2007; Sherrill et al., 2013). Future studies are necessary to examine potential age differences in METH sensitivity more directly. For example, breakpoints under a PR schedule for different unit doses could be measured before and after escalation. If reduced sensitivity significantly contributes to escalation of METH intake, a downward shift in the dose-response curve for breakpoints post-escalation, as well as adolescents reaching lower breakpoints than adults, would be the likely outcomes. On the other hand, if sensitization to the reinforcing effects of METH contributes to escalation of intake, an upward shift in the dose-response curve for breakpoints post-escalation would be the most likely outcome.

An alternative explanation for the age-of-onset difference in escalation of METH intake is that adolescent-onset rats transition more readily from controlled to habitual or compulsive drug-taking. One way to investigate this idea is to measure responding when the drug is no longer available, for example in a signaled non-availability paradigm, as continued drug-seeking may indicate that the response has become habitual or compulsive. Previous work using cocaine and food as reinforcers has shown that adolescent rats of both sexes responded more during signaled non-availability compared to adult rats (Anker et al., 2011), supporting the notion that adolescent rats may develop habitual/compulsive drug-seeking more readily than adult rats. Furthermore, age differences in compulsive drug-taking have also been assessed when drug-taking behavior becomes punished to model continued drug use despite negative consequences. For example, Holtz and Carroll (2015) trained adolescent and adult female rats to self-administer cocaine under ShA conditions for 10 days before adding an aversive agent, histamine, to the cocaine solution. With histamine present, adult rats self-administered less cocaine than adolescents. Taken together, these findings suggest that adolescent rats exhibit greater compulsive drug-taking compared to adults, which may partly explain the greater escalation of METH intake in adolescents seen in the current study.

Both sex and age-of-onset differences were evident with self-administration of the non-drug reinforcer, saccharin. During ShA and LgA sessions, females earned more saccharin than males when normalized to body weight. In stark contrast to METH escalation during LgA, rats rapidly reduced their intake of saccharin when given access for 6 h, suggesting that escalation of intake may be specific to drug reinforcers. We chose non-caloric saccharin as our non-drug reinforcer in order to control for the influence of differing metabolic needs between adults and rapidly growing adolescents. However, when given longer access to saccharin, rats may rapidly learn the lack of nutritive effects of the reinforcer (i.e., that saccharin is non-caloric) and decrease their intake accordingly. This notion is consistent with prior studies in adult male mice (Sclafani et al., 2015). In this study, a non-caloric saccharin and sucralose mixture was preferred to caloric sucrose or glucose solutions in brief (1 min) access choice tests, but this preference switched across time when mice were given 2 days of 24-h access to these solutions. These results suggest that the post-oral nutritive effects, rather than palatability, influenced sweetener intake across time (Sclafani et al., 2015). Despite the overall reduction in saccharin intake across time in the current study, adolescent-onset and female rats maintained higher saccharin intake than their adult-onset and male counterparts. These group differences are consistent with a body of literature that has demonstrated hypersensitivity to non-drug rewards in adolescents and females (Friemel et al., 2010; Hammerslag and Gulley, 2014; Marshall et al., 2017; Hankosky et al., 2018; Westbrook et al., 2018).

During the period when rats were engaged in daily self-administration sessions, all animals gained weight as expected for the Sprague-Dawley strain. However, adolescent-onset male rats self-administering METH exhibited suppressed weight gain starting during the second week of LgA, which coincided with escalation of their METH intake. METH is known to have appetite-suppressing effects, and human METH abusers often experience weight loss (Hawks et al., 1969). In rodents, adult male rats that are given extended access to METH at the same dose used in the current study (0.1 mg/kg/inf) have been reported to lose weight (Krasnova et al., 2010). We did not find significant weight loss in our adult-onset males, however, and this discrepancy may be due to differences in session duration. Our study used the more common 6-h duration for 14 daily sessions, whereas Krasnova and colleagues used longer duration access over fewer days (14-h for 8 days). Nonetheless, we did find METH-induced suppression of weight gain in our adolescent-onset male rats. Under normal conditions, adolescent males gain weight at the highest rate compared to the other groups, which may make it easier to detect more subtle changes in weight gain from METH self-administration in our LgA paradigm. Notably, this suppression of body weight gain is not long-lasting. About 1 week after cessation of self-administration, body weights of adolescent-onset METH rats rebounded to reach similar values as adolescent-onset rats that self-administered saccharin.

Approximately one-two weeks after their last self-administration session, rats were tested on recognition memory tasks. We saw no group differences in exploration time during the study phase for either the NOR or OiP tasks, suggesting that self-administration history did not influence exploratory behavior in general. In tests of recognition memory, we found no evidence for sex differences, regardless of self-administration history. This lack of a sex effect is consistent with a previous report in adult rats with a history of METH self-administration during adulthood (Reichel et al., 2012a). In line with our hypothesis, all rats recognized the familiar object and spent more time exploring the novel object in the NOR task. Our findings are consistent with previous studies reporting no NOR deficits after 1 week of abstinence from non-contingent METH administration during adolescence (North et al., 2013) or adulthood (Melo et al., 2012) in male mice. However, Reichel and colleagues have shown NOR impairments in adult rats after LgA METH self-administration (Reichel et al., 2011, 2012a,b). Although the design for our current study was based on the work from Reichel and colleagues, our study did differ from these studies in two ways. Our study used Sprague-Dawley rats, whereas Reichel and colleagues used Long-Evans in their experiments. There may be strain differences in susceptibility to cognitive deficits following METH exposure. Another notable difference is that our rats were group housed, while those in Reichel and colleagues’ studies were individually housed. We chose to group house our rats in order to reduce any potential age differences in the social isolation stress, as adolescence is a time of increased social interaction relative to adulthood (Douglas et al., 2004). Moreover, social housing has been reported to enhance novel object recognition memory in mice of both sexes compared to individual housing (Liu et al., 2019). Thus, our approach of minimizing housing-induced stress may have helped protect against METH-induced deficits in NOR.

We found that OiP memory was not as robust as NOR memory in our naïve control rats, which is consistent with previous literature (Dix and Aggleton, 1999). The naïve control group did display significant preference for the novel change in OiP during the first min of the test phase, demonstrating that this group was able to recognize that the objects switched places. However, this group appeared to habituate rapidly to this change across the three-min test session, resulting in the total test phase exploration ratio not being significantly different from chance performance. This rapid habituation was not apparent in the groups with a history of self-administration. In rats that self-administered SACC beginning during adolescence or adulthood, we did not find evidence for impaired OiP memory. This finding is in contrast to a previous report showing deficits in OiP and associated neural changes in the mPFC and hippocampus in adulthood following adolescent (P28-56) sucrose access (Reichelt et al., 2015). This discrepancy may be due to differences in the non-drug rewards used, such as their nutritive qualities (caloric vs. non-caloric) or the testing paradigm (free access vs. operant self-administration). Consistent with our predictions, an age-of-onset dependent deficit was present in OiP recognition in rats with a history of METH use, although this effect was fairly modest. Rats that previously self-administered METH beginning in adolescence displayed significantly lower exploration ratios than rats that began in adulthood. Furthermore, adolescent-onset METH self-administering rats failed to demonstrate significant recognition of the novel change in any of the three min in the test phase, while adult-onset METH self-administering rats recognized the change by the first min when the novelty salience of position change is at its height. However, it is important to note that neither age-of-onset group of rats with a history of METH use significantly differed in OiP performance from the naïve control rats.

The PFC is required for OiP recognition as lesions have been shown to selectively impair memory in this task but not in tests of NOR memory (Barker et al., 2007). Moreover, D_1_Rs mediate OiP memory as pharmacological blockade of D_1_Rs in the PFC results in task-specific deficits in OiP recognition that do not occur with D_1_R blockade in the perirhinal cortex or hippocampus (Savalli et al., 2015). The specific deficit in OiP in adolescent-onset compared to adult-onset METH self-administering rats suggests that METH use during adolescence may impact the PFC, hippocampus, or both brain regions (Barker and Warburton, 2011), but not the perirhinal cortex or general sensory, motor, or motivational capacities because NOR memory was intact. Although only a modest effect, our recognition memory findings are consistent with our hypothesis that METH use during this period adversely impacts developmental events in the PFC, although more work is needed to determine whether the hippocampus is also impacted.

We measured D_1_R and NMDAR subunit protein expression in the PFC and NA following three weeks of abstinence from METH or saccharin self-administration. Our findings of no significant group differences for any of our proteins of interest in either brain region are in contrast to previous reports from our lab (Kang et al., 2016b) and others (Caffino et al., 2018). Notably, the previous studies used non-contingent (i.e., experimenter-administered) exposure to psychostimulants. Voluntary versus involuntary drug administration has been reported to differentially influence the brain and behavior, including the extent of the drug’s effects on dendritic spine morphology in the PFC (Radley et al., 2015). It may be the case that the added stress of non-contingent drug injections contributes to the drug-induced neuroadaptations following adolescent drug exposure reported in previous studies (Kang et al., 2016b; Caffino et al., 2018).

Our inability to find a change in GluN1 NMDAR subunit expression in the mPFC following LgA METH self-administration is consistent with previous studies in adult male rats that self-administered cocaine (Ben-shahar et al., 2007) or METH (Reichel et al., 2014; Mishra et al., 2017) under LgA conditions. However, the findings for GluN2B surface expression in the mPFC were mixed with one study reporting no change (Reichel et al., 2014) and the other an increase (Mishra et al., 2017). These discrepancies may be explained by the region(s) of mPFC that were assessed. The regions of mPFC collected by Mishra and colleagues (2017) were not specified, whereas the study by Reichel and colleagues (2014) and the current report analyzed the prelimbic and infralimbic subregions of the mPFC together in a single homogenate. We chose not to separately analyze these subregions because of our lab’s previous work showing no difference between them following adolescent or adult AMPH exposure (Kang et al., 2016a). While combining the two regions does not explain our lack of differences in D_1_R expression, it may account for our null findings for GluN2B as Caffino and colleagues (2018) reported opposing directions of effect of cocaine injections during adolescence on phosphorylated GluN2B expression in the prelimbic versus infralimbic subregions. Another consideration is that our rats underwent their final recognition memory test 1 week prior to tissue collection, which has the potential to influence our findings. However, in light of a study showing GluN2B expression in the PFC was not significantly altered following training or testing in recognition memory (Cercato et al., 2017), it is likely the case that our measure of protein expression would be influenced by drug-induced adaptations and not task-related learning or performance. It is possible that the functional trajectories of D_1_R and/or GluN2B in the PFC are disrupted without a concomitant change in protein expression. In support of this notion, our lab previously reported long-lasting reductions in sensitivity to D_1_R agonists in the PFC after non-contingent AMPH exposure during adolescence (Kang et al., 2016a).

In conclusion, our investigation into METH-induced cognitive dysfunction and neuroadaptations in D_1_Rs and NMDARs following contingent METH use revealed age-of-onset and sex differences in escalation of METH intake. In addition to these differences in drug-taking patterns, the current study provides some evidence for modest disruptions in PFC-dependent cognitive functions when METH use begins during adolescence; however, there appears to be no long-lasting impacts on the ontogeny of D_1_R and GluN2B expression in the PFC. Overall, these findings suggest that adolescent-onset users may develop problematic drug-taking patterns more rapidly and may be more likely than adult-onset users to experience cognitive dysfunction in early abstinence. Both of these factors could contribute to higher drop-out rates from treatment programs (McKellar et al., 2006), greater relapse rates (Simon et al., 2004; Poudel and Gautam, 2017), and worse treatment outcomes (Hillhouse et al., 2007). Furthermore, adolescent-onset users may benefit more from treatment programs that focus on improving cognitive function during early abstinence.

## Acknowledgements

The authors thank Jessica Alvarez (supported by an NSF/REU award), Erika Carlson, Qingrou (JoJo) Gu, Kate Hamblen, Kristen Hughes, Adrianna Jelen, Shawn Kurian, Karen Lai, Jacob O’Russa, Sarah Rahman, Brittany Rhed, Tugba Serbest, and Ashley Wehrheim for excellent technical assistance.

## Funding

This work was supported by funding from the National Institutes of Health (DA 029815), the University of Illinois Campus Research Board, and an NSF REU award (1559908/1559929).

## Notes

#### Summary of Updates

Revised version following response to reviewer critiques

