## Supplemental Tables for "Extended access self-administration of methamphetamine is associated with age- and sex-dependent differences in drug taking behavior and recognition memory deficits in rats"

**Supplementary Material**

**Supplementary Table 1.** Total test phase exploration ratios compared against chance (0.5) in recognition memory tasks.

| **Task** | **SA History** | **Age-of-onset** | **df** | **t** | **FDR-adjusted p-value** |
| --- | --- | --- | --- | --- | --- |
| OR | METH | Adolescent | 25 | 6.99 | **0.0003*** |
|  |  | Adult | 32 | 9.07 | **0.0003*** |
|  | SACC | Adolescent | 27 | 5.21 | **0.0003*** |
|  |  | Adult | 29 | 9.69 | **0.0003*** |
|  | Naïve | Adult | 23 | 5.29 | **0.0003*** |
| OiP | METH | Adolescent | 25 | -0.26 | 0.8597 |
|  |  | Adult | 32 | 2.91 | **0.0118*** |
|  | SACC | Adolescent | 27 | 2.43 | **0.0365*** |
|  |  | Adult | 29 | 0.51 | 0.7688 |
|  | Naïve | Adult | 23 | 0.21 | 0.8791 |

***indicates significant p-values after adjusting for multiple comparisons**

**Supplementary Table 2.** Minute breakdown of test phase exploration ratios compared against chance (0.5) in recognition memory tasks.

| **Task** | **SA History** | **Age-of-onset** | **Minute** | **df** | **t** | **FDR-adjusted p-value** |
| --- | --- | --- | --- | --- | --- | --- |
| OR | METH | Adolescent | 1 | 25 | 6.33 | **0.00029*** |
|  |  |  | 2 | 25 | 3.46 | **0.00380*** |
|  |  |  | 3 | 25 | 3.89 | **0.00165*** |
|  |  | Adult | 1 | 32 | 9.94 | **0.00029*** |
|  |  |  | 2 | 32 | 5.99 | **0.00029*** |
|  |  |  | 3 | 32 | 4.19 | **0.00053*** |
|  | SACC | Adolescent | 1 | 27 | 7.14 | **0.00029*** |
|  |  |  | 2 | 27 | 5.06 | **0.00029*** |
|  |  |  | 3 | 27 | 1.00 | 0.43293 |
|  |  | Adult | 1 | 29 | 8.81 | **0.00029*** |
|  |  |  | 2 | 29 | 5.92 | **0.00029*** |
|  |  |  | 3 | 29 | 5.63 | **0.00029*** |
|  | Naïve | Adult | 1 | 23 | 7.88 | **0.00029*** |
|  |  |  | 2 | 23 | 2.27 | 0.05092 |
|  |  |  | 3 | 23 | 3.64 | **0.00295*** |
| OiP | METH | Adolescent | 1 | 25 | 1.13 | 0.37090 |
|  |  |  | 2 | 25 | -1.49 | 0.21300 |
|  |  |  | 3 | 25 | 0.03 | 0.97650 |
|  |  | Adult | 1 | 32 | 2.72 | **0.01826*** |
|  |  |  | 2 | 32 | 0.47 | 0.77212 |
|  |  |  | 3 | 32 | 3.11 | **0.00743*** |
|  | SACC | Adolescent | 1 | 27 | 0.29 | 0.85967 |
|  |  |  | 2 | 27 | 2.32 | **0.04528*** |
|  |  |  | 3 | 27 | 0.42 | 0.77863 |
|  |  | Adult | 1 | 29 | 3.60 | **0.00267*** |
|  |  |  | 2 | 29 | 0.13 | 0.92359 |
|  |  |  | 3 | 29 | -0.55 | 0.75910 |
|  | Naïve | Adult | 1 | 23 | 4.03 | **0.00125*** |
|  |  |  | 2 | 23 | 0.45 | 0.77212 |
|  |  |  | 3 | 23 | -1.81 | 0.12444 |

***indicates significant p-values after adjusting for multiple comparisons**
